# Weaver ants regulate the rate of prey delivery during collective vertical transport

**DOI:** 10.1101/2022.06.22.497253

**Authors:** Andrew T. Burchill, Theodore P. Pavlic, Stephen C. Pratt, Chris R. Reid

## Abstract

The collective transport of massive food items by ant teams is a striking example of biological cooperation, but it remains unclear how these decentralized teams coordinate to overcome the various challenges associated with transport. Previous research has focused on transport across horizontal surfaces and very shallow inclines, disregarding the complexity of natural foraging environments. In the ant *Oecophylla smaragdina*, prey are routinely carried up tree trunks to arboreal nests. Using this species, we induce collective transport over a variety of angled surfaces with varying prey masses to investigate how ants respond to inclines. We found that weight and incline pose qualitatively different challenges during transport. Prey were carried over vertical inclines *faster* than across horizontal surfaces even though inclines were associated with longer routes and a higher probability of dropping the load. This additional speed is associated with more transporters being allocated to vertical loads and not from the persistence of individual ants. Ant teams also regulated a stable “prey delivery rate” (rate of return per transporter) across all treatments. Our proposed constrained optimization model explains these results: prey intake rate at the colony level is maximized when the allocation of transporters yields a similar prey delivery rate across loads.

## Introduction

Coordination allows groups of organisms to perform tasks that are insurmountable for individuals. For example, in some species of ants, workers will cooperatively transport food items that are too large for a single forager to carry. Cooperative transport requires that teams contend with many different challenges; loads of highly variable size, shape, and weight must be retrieved and carried around obstacles and over complex terrain (McCreery and Breed 2014). Team members must take on appropriate roles and be prepared to change their behavior rapidly as conditions change (for example, the load rotates or slips, or other ants join or leave the group). The team must solve these challenges without direction by a well-informed leader, instead relying on decentralized mechanisms of coordination that remain poorly understood.

Insight into the ants’ behavior can be gained by examining how their transport performance scales with load size. Some group-raiding species (marauder ants and army ants) become more efficient at larger load sizes (Franks 1986, Moffett 1988a, Franks et al. 1999). This “superefficiency” can be quantified by the prey delivery rate (PDR). PDR is calculated as the load’s mass multiplied by its velocity and divided by the number of active transporters; that is, PDR is an index of the average individual’s contribution to the mass flux in a cooperative-transport team (Buffin and Pratt 2016). Larger PDR values indicate that fewer ants are needed to carry the same object at the same speed. For army ants, at least, rising PDR with load size is attributable to their distinctive transport behavior in which porters straddle loads and face in the same direction, resulting in reduced conflict and rotational forces (Franks et al. 2001). For most ants, however, PDR declines with load size (Buffin and Pratt 2016), and teams carrying larger loads move more slowly, involve more transporters, and may take a more sinuous path back to the nest (Detrain 1990, Cerdá et al. 2009, Buffin and Pratt 2016, McCreery et al. 2019). These deficiencies may reflect constraints imposed by how they carry loads and coordinate with one another, but the ways in which load size affects transport behavior remain poorly understood.

Even less attention has been paid to how the transport environment affects group performance. Most research has been conducted over artificial, featureless surfaces (Moffett 1988b, Czaczkes et al. 2011, McCreery 2017, Buffin et al. 2018). Several studies have examined how ant teams navigate around discrete obstacles (McCreery et al. 2016, Gelblum et al. 2020), but these investigations have still been restricted to flat, horizontal planes (except see (Qin et al. 2019), which focuses on how ants disassemble artificial loads on trees). In nature, ants are often seen transporting through leaf litter, across vines, and up tree trunks. Transport teams in some species regularly scale sheer vertical surfaces (Wojtusiak et al. 1995) and even carry objects upside-down along the underside of horizontal surfaces. Such inclines present several unique challenges – including supporting the load’s weight against gravity and overcoming inevitable mistakes that can reverse progress when not quickly corrected. Many ant species are very capable of cooperative transport despite these challenges, and yet the coordinating mechanisms that allow them to do this remain unknown.

The differences between vertical and horizontal collective transport as well as the implications of said differences on ant behavior are not immediately clear. Previous studies present somewhat conflicting evidence about possible energetic costs. Among leaf-cutter ants, individual transport while ascending vertical surfaces seems to be more energy intensive than horizontal transport (Lewis et al. 2008), but in unladen carpenter ants, vertical locomotion is not more metabolically costly (Lipp et al. 2005). Additionally, Endlein and Federle (2015) have shown that weaver ants employ different biomechanics when walking over various inclines, using either their arolia or tarsal claws. However, these studies only consider individual ants, and it is not clear whether these findings would translate to individuals coordinating motion in transport groups. Furthermore, vertical transport not only presents an energetic challenge but also a logistic one as mistakes (e.g., dropping loads) that would be minor on horizontal surfaces can be catastrophic on steep inclines. These mistakes may occur more frequently with teams as the objects they carry could be significantly heavier than any one ant (or even subset of the team) can carry. Moreover, other challenges such as increasing load weight may be exacerbated on vertical surfaces.

To better understand how ants that are effective at cooperative transport overcome the likely challenges unique to steep inclines, we focus our study on green weaver ants, *Oecophylla smaragdina*, a highly polydomous, arboreal-nesting species with very large colonies. *Oecophylla* are voracious hunters that can scavenge upon small lizards, mice, and birds (Wojtusiak et al. 1995). They forage both among the trees and along the ground, and so they must regularly engage in both vertical and horizontal transport of these relatively large prey items. Given their ability to transport across inclines and collect prey of vastly different masses, *Oecophylla smaragdina* represents an ideal model of how transport efficiency is affected by the challenges posed by load weight, incline, and their potential interaction. By presenting multiple challenges simultaneously, we can also better tease apart how the various subcomponent measures of group performance interact. Additionally, we can test whether vertical collective transport poses a logistical challenge: under what circumstances might slipping or falling backwards occur, and do the ants employ mechanisms to mitigate this risk? To investigate these questions, we offered field colonies loads of various weights across different inclines while capturing video of cooperative transport bouts and measuring team performance. We then quantified the effects of incline and weight, as well as their interaction, on multiple aspects of transport performance.

## Methods

### Overview

We coerced weaver ant colonies to establish trails over experimental platforms with angles of inclination that were either 0, 45, or 90° from horizontal. We presented these colonies with prey items that differed in mass, with treatments of 0.25, 1.25, or 2.25 g (plus minus some SD). We video recorded the platforms as ant teams collectively transported the loads toward their nests. From these video data, we tracked the position of the load and the number of ants engaged with the load over time. We also measured the duration for which a subset of randomly chosen individual ants remained engaged with the load. We then calculated multiple metrics of transport performance, including speed, straightness, and prey delivery rate, and we examined the effects of incline and load weight on these factors.

### Field site

Experiments were conducted during midday (10:00 – 15:00) between October 16th and November 5th, 2018 in Kirwan, a suburb of Townsville in Queensland, Australia. We studied three polydomous colonies of *Oecophylla smaragdina*, with each possessing nests in multiple nearby trees (although data from one colony was excluded from final analysis, see below). We determined colony identities by manually swapping workers between trees and monitoring them for aggressive interactions. The activity level of each nest--determined by visually assessing the rate of foraging trail traffic--was found to be approximately the same.

### Experimental details

Prior to any experimental manipulation, we chose trees with active ant trails within the territory of each colony. Adapting methods from Romeu□Dalmau et al. (2010), we wrapped the lower portion of these trunks in plastic sheets dusted with talcum powder. This disrupted the existing trails, as ants would not cross the coated plastic. We then attached cardboard bridges across these obstacles, along which the ants’ trails were quickly reestablished.

The following day, we placed the 65 cm by 35 cm experimental platforms (made from cardboard layered over particle board panels) at the base of these trees, at inclines of either 0°, 45°, or 90° (n = 36, 36, and 34 respectively) from horizontal (Fig 1). We determined the exact angles using spirit levels and the iPhone 6s gyroscope. The cardboard bridges were adjusted so that the active trails would continue down the center of the platform and so that ants could not cross the plastic barrier without walking over the platform. This setup coerced ants into forming trails over the platforms, thus minimizing recruitment latency. We used fresh cardboard platforms for each colony, each day. To control for differences in the initial “ramp-up” phases of recruitment and foraging, we placed previously freeze-killed crickets (∼0.25g) at the far end of each platform before the experiment began. Experiments did not commence until the initial cricket had been transported to the nest and the trail had been thoroughly established.

**Figure 1:**
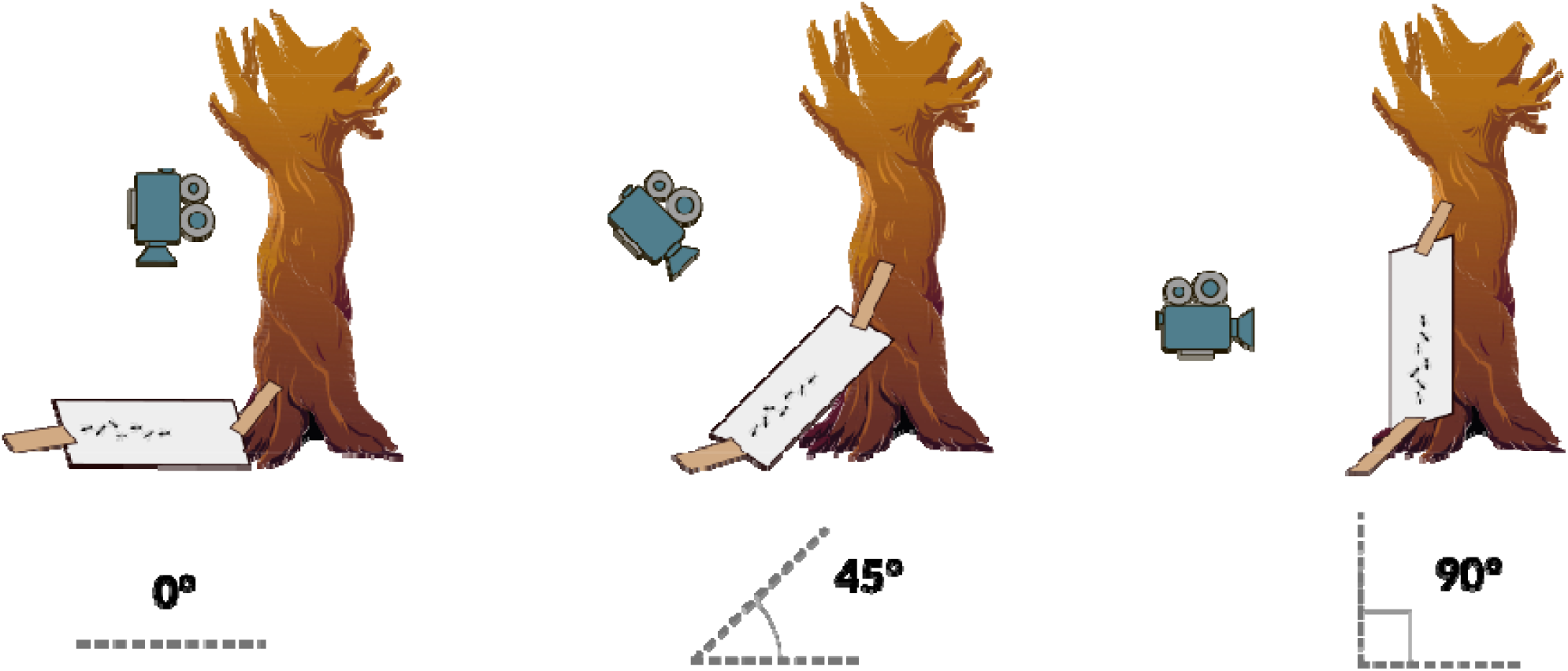
Cartoon illustrating the placement of experimental platforms and the constant filming angle relative to the platforms.

We recorded transport in Cinematic 4K video with Panasonic GH4 cameras aimed and positioned orthogonally to the incline of the platforms at a distance of approximately one meter. This rendered similar “bird’s-eye views” regardless of platform angle. Our prey items were freeze-killed crickets of similar size, thawed and delimbed, and either unadorned or with one or two lead weights attached via hot glue as standard loads, similar to Lioni (2000). The weights of these light, medium, and heavy weight classes were approximately 0.25g, 1.25g, and 2.25g respectively (plus minus some SD). We recorded the exact weight of each load before presenting it to the ants.

For each colony, we selected three inhabited trees with nests and active foraging trails for our experiment. To remove the possible confounding effect of a particular tree, we presented each tree with each angled platform over three subsequent days. Each day of recording, we presented all trees with all three prey weight classes, randomly ordered, twice. After each load was transported to the end of the platform, we removed the attached ants from the load and the experiment and then immediately returned the load for a second run. After its second run, the load was replaced with a new load from a different weight class, and the process was repeated. Unfortunately, unexpected water damage warped our experimental platforms during two days of filming, so we did not include data from the affected colony in the final analysis.

### Data extraction

We extracted load trajectories and data on individual transporters from the videos. We converted the transport videos to five frames per second and imported them into Fiji v1.52 (Schindelin et al. 2012), where the position of the cricket load’s head was manually tracked every 60 frames (12 seconds). To control for differences in initial recruitment dynamics, we only analyzed data collected after the load had moved 5 cm. We considered runs completed once the load reached the end of the platform nearest the nest.

#### Average displacement speed and average instantaneous speed

From the load trajectory data, we calculated two alternative measures of transportation speed. “Average displacement speed” is the straight-line distance between the load’s initial and final positions, divided by the total time it took to move between them. The “average instantaneous speed” represents the moment-to-moment magnitude of the load’s velocity in any direction, averaged over the entire run. Average displacement speed and average instantaneous speed would be identical if a load were to move directly toward its ending position, but average displacement speed would be lower than average instantaneous speed if a load were to take a more circuitous path.

#### “Backtracking”

We occasionally observed loads slipping backwards and falling, and we wanted to determine what factors led to this behavior during runs. Due to the difficulty in accurately discriminating between slips caused by gravity and general backwards movement, we use the equivocal term “backtracking” to refer to both. We calculated the amount of “backtracking” as the sum of distances covered when teams were moving further from the nest (and thus towards the ground), normalized by the total start-to-end distance the load moved during its run.

#### Straightness of the load’s path

We calculated path straightness by dividing the distance between the load’s initial and final positions by the total distance it traveled. To avoid conflating general path straightness with backtracking events, we subtracted path segments that were classified as periods of backtracking from the total distance traveled.

#### Group size and individual persistence

We recorded the number of transporters involved with each load over time. We considered an individual ant engaged with the load when the ant’s mandibles were visually overlapping with the load for more than two consecutive video frames. We then manually recorded the number of ants engaged with the load every 120 frames (24 seconds) or, if the video was particularly long, a multiple of 120 frames that resulted in at least five records. Because transporter numbers tended to increase initially before plateauing, we used the median number of transporters over time for each run.

To determine how long individual ants remained engaged with the load, we first chose a random frame between one quarter and three quarters of the way through each video. A randomly chosen individual engaged with the load at this time-point was then manually followed frame-by-frame until the individual let go of the load and remained disengaged for two consecutive frames. This duration was recorded for n=194 ants.

### Statistical analysis

All statistical analyses were completed using R version 3.6.1 (R Core Team 2019).

#### Question 1: How do challenges affect transport efficiency?

The efficiency or ability of ant teams to collect massive food objects is often measured by prey delivery rate (or PDR) in ant collective transport studies. PDR is usually defined as the mass of the load being retrieved, multiplied by its average displacement speed, and divided by the number of ants transporting it. In large ant colonies with hundreds to thousands of foragers simultaneously retrieving prey items, the net intake rate per ant likely represents a highly ecologically relevant metric, and it allows for comparison across species as well as across load and group sizes.

Previous work has shown that the challenge of increasing load weight often decreases transport efficiency and that larger team sizes are associated with this decrease (Buffin and Pratt 2016). Thus, we examined the effect of three potential predictor variables (incline, load weight, and the median group size) on the PDR using linear mixed-effect models with the random effects of date nested within colony id. We applied a square root transform to the PDR values to make them more normally distributed. We compared the full model containing all three predictors and their interactions with all simpler sub-models. Additionally, because we did not find any predictor to have a significant effect on PDR in any of the models, we also used a multi-model averaging approach to quantify possible effect sizes, employing the methods used in Camacho et al (2018).

#### Question 2: How do more granular measures of group performance such as transportation speed, path straightness, degree of backtracking, and group size respond to changes in substrate incline and load weight?

We used general linear mixed-effect models to analyze the effects of incline and load weight on average displacement speed, average instantaneous speed, the straightness of the path taken by the load, and the median number of ants engaged with the load over the run. For each dependent variable, we used stepwise regression and likelihood ratio tests to determine the model which best matched the observations.

We log-transformed velocity and speed data to improve normality and reduce heteroscedasticity. After transformation, the Q–Q plots showed very slight deviations from normality, but given our relatively large sample size (n=106), the increased homogeneity of variance among our residuals, and the fact that Gaussian models are fairly robust to violations of normality (Knief and Forstmeier 2021), we felt confident using the log-transformed speed and velocity data in the linear mixed-effect models. Given that path straightness as a value is bounded between 0 and 1, we applied an arcsine square root transform to the straightness data before analysis (Sokal and Rohlf 1995).

To analyze how load weight and incline lead to backtracking during transport, we used a hurdle mixed model from the R package *GLMMadaptive*. Specifically, a hurdle (or “two part”) model uses a standard linear mixed model for the log-transformed backtracking distance of the non-zero responses, as well as a logistic regression for whether any backtracking occurred at all. We included both incline and load weight as possible predictor variables for the linear and logistic components, and we used colony identity as a random effect in both components as well. We used stepwise regression and likelihood ratio tests to determine the model which best describes the observed results.

#### Question 3: How do individual persistence and group size affect instantaneous movement speed?

After finding a statistically significant, positive association between steeper inclines and increased instantaneous movement speed, we used our data to test the validity of several hypotheses that could explain this counter-intuitive result. Specifically, we tested the (not mutually exclusive) hypotheses that the “quality” of individual transporters and/or their quantity were responsible for this association.

##### Survival analysis

We examined how long individual ants remained locally engaged with loads via survival analysis. We used a mixed-effect Cox regression from the R package *coxme*, with incline and load weight as our predictor variables, and included the random effects of trial nested within date within colony identity. We used stepwise regression to compare a full model with both predictors and the set of simpler models.

##### Mediation analysis

To explore the underlying causal relationship between steeper slopes and load speed, we also employed a mediation model. Mediation analysis, a common tool in the social sciences, tests whether a third variable can explain (or mediate) the apparent effect of an independent variable on a dependent variable. We wanted to test whether incline’s effect on load speed was mediated through transporter team size, which was larger on steeper slopes.

To model the relationship of group size, load weight, and incline on average instantaneous load speed, we ran a full linear regression using all three predictors and then all simpler sub-models, determining the best predictor variable(s) based on Bayesian information criterion (BIC). We tested the significance of this indirect effect using quasi-Bayesian procedures from the R package *mediation*. Unstandardized indirect effects were computed for the 1000 simulation samples.

## Results

### Prey delivery rate

We found that none of our predictors—in any of our sub-models—had a significant effect on prey delivery rate (PDR) (Fig 2A). Furthermore, the potential effect size of each of our predictors is less than the effect on PDR of adding a single, theoretical lazy ant to the largest transport teams carrying the heaviest loads. That is, while transporting the heaviest loads, ants would continuously be engaging and disengaging from the load. We can calculate the maximum decrease in PDR that would result from the addition of a single, theoretical lazy ant to these already large (15+ ants) transport teams. This unhelpful individual’s presence would decrease the PDR (which is calculated per capita) by 0.00161—, which is larger than the model-averaged effect sizes for *any* of our predictors. For example, the predicted difference in PDR between loads carried over vertical and horizontal surfaces is only 0.00154—. So load weight, incline, and the number of transporters have, at most, a biologically negligible effect on PDR, with such an effect being practically equivalent to zero.

**Figure 2:**
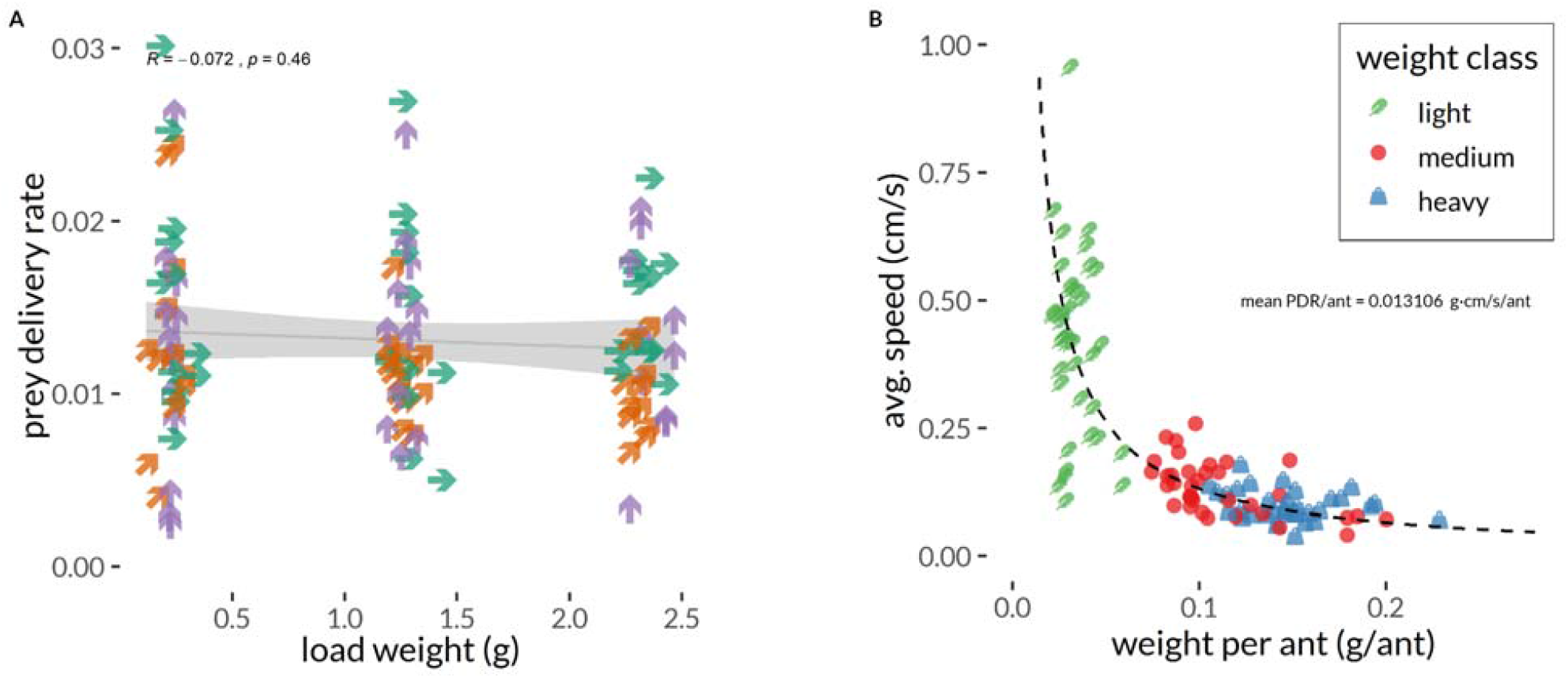
Prey delivery rate remains constant across all treatments. (A) Neither load weight nor incline (indicated by direction and color of the points) had a biologically meaningful effect on the rate of return per transporter. (B) Load speed and the weight carried per ant have a tight, inversely proportional relationship that suggests a possible trade-off between the two factors. Light loads are transported quickly but contribute little prey mass to the colony, while heavy loads yield more mass per transporter yet take longer.

### Measures of group performance

#### Average displacement speed

Although increasing load weight decreased the average displacement speed of transportation (* = -0.33, *p <* 0.001, *t*_*99*_ = -12.98, *R*^*2*^ *=* 0.576), the angle of inclination did not have a significant effect (Fig 3). Likewise, we found no significant interaction between load weight and the angle of the experimental platforms on the average displacement speed. On average, loads were transported from the beginning to the end of our platforms at velocities of 0.420 cm/s, 0.125 cm/s, and 0.0876 cm/s for the light, medium, and heavy loads respectively.

**Figure 3:**
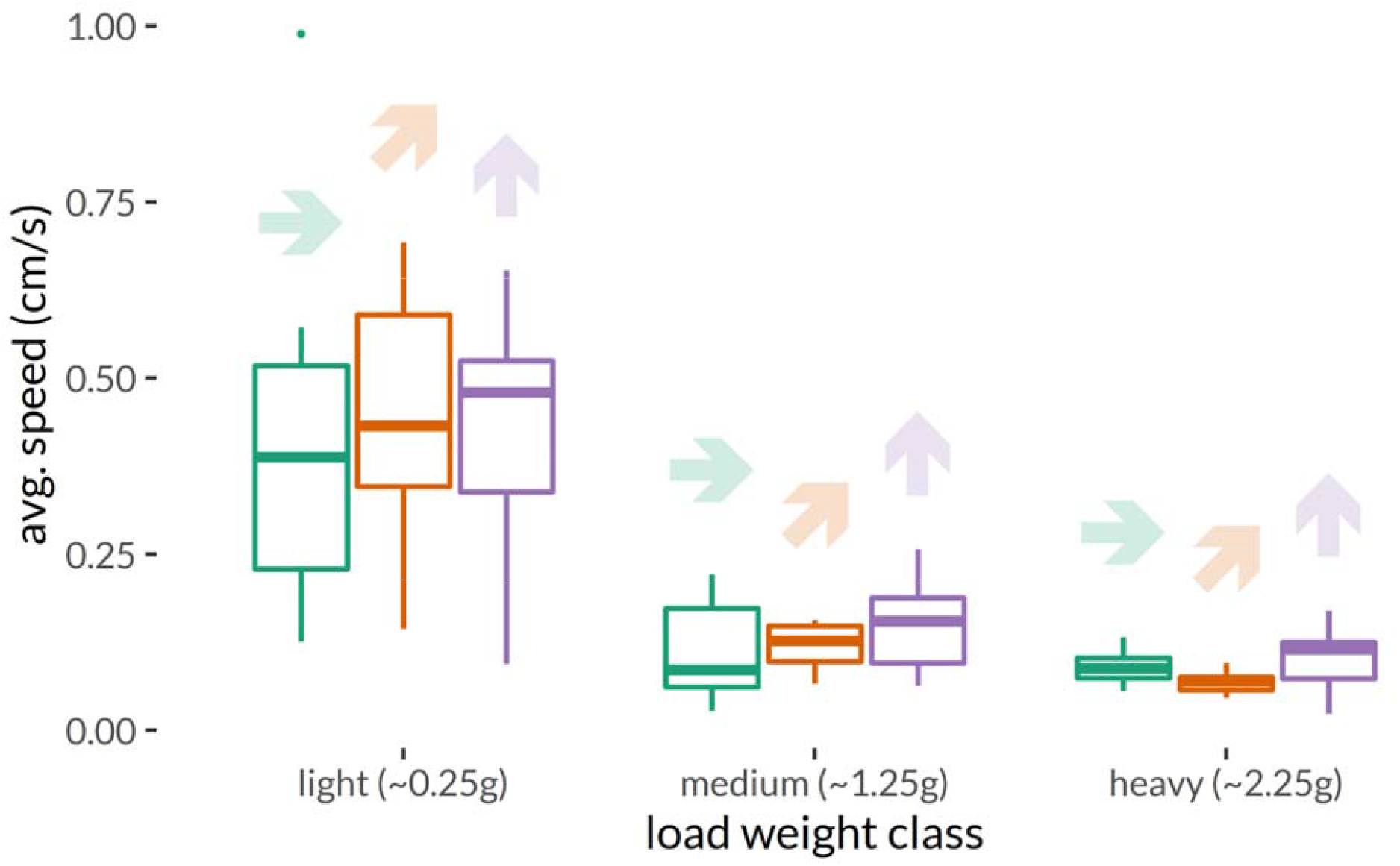
The average displacement speed of loads as a function of load weight and incline. Speed decreases with heavier loads but not with steeper inclines.

#### Group size

Both steeper slopes (* = 0.30, *p <* 0.001, *t*_*98*_ = 6.89) and heavier loads (* = 0.82, *p <* 0.001, *t*_*98*_ = 18.6) increased the median number of transporters engaged with a load, although there was not a significant interaction between the two factors (Fig 4. *R*^*2*^ *=* 0.746). All things being equal, group size would be expected to increase from ∼10 ants on a horizontal platform to ∼13 on a vertical surface and from ∼7 ants on a 0.25 gram load to ∼17 on a 2.25 gram load.

**Figure 4:**
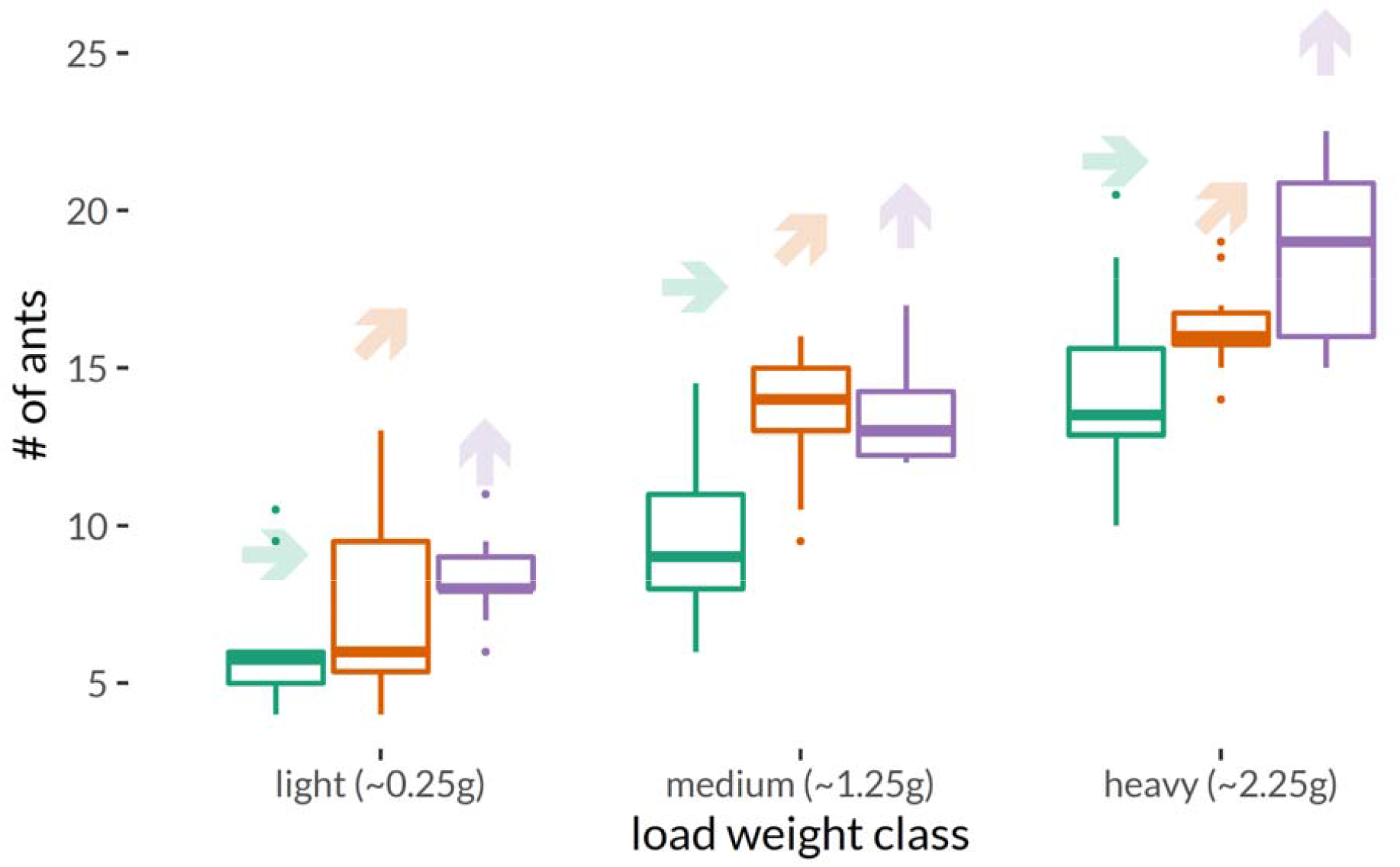
The median number of transporters engaged with a load as a function of load weight and incline. Significantly more ants are involved with heavier loads and on steeper inclines.

#### Backtracking and slipping

Backtracking was more likely to happen when loads were heavier (*_zi_ = -1.45, *p <* 0.001, *z* = -4.52), and the distance backtracked was greater on steeper slopes (* = 0.67, *p <* 0.001, *z* = 3.38). The weight of the load predicted the likelihood of whether backtracking occurred during transportation, with lighter loads being less likely to backtrack (*R*^*2*^ *=* 0.187). However, the distance backtracked was significantly affected only by the inclination of the experimental platform. Steeper slopes resulted in longer distances backtracked (Fig 5).

**Figure 5:**
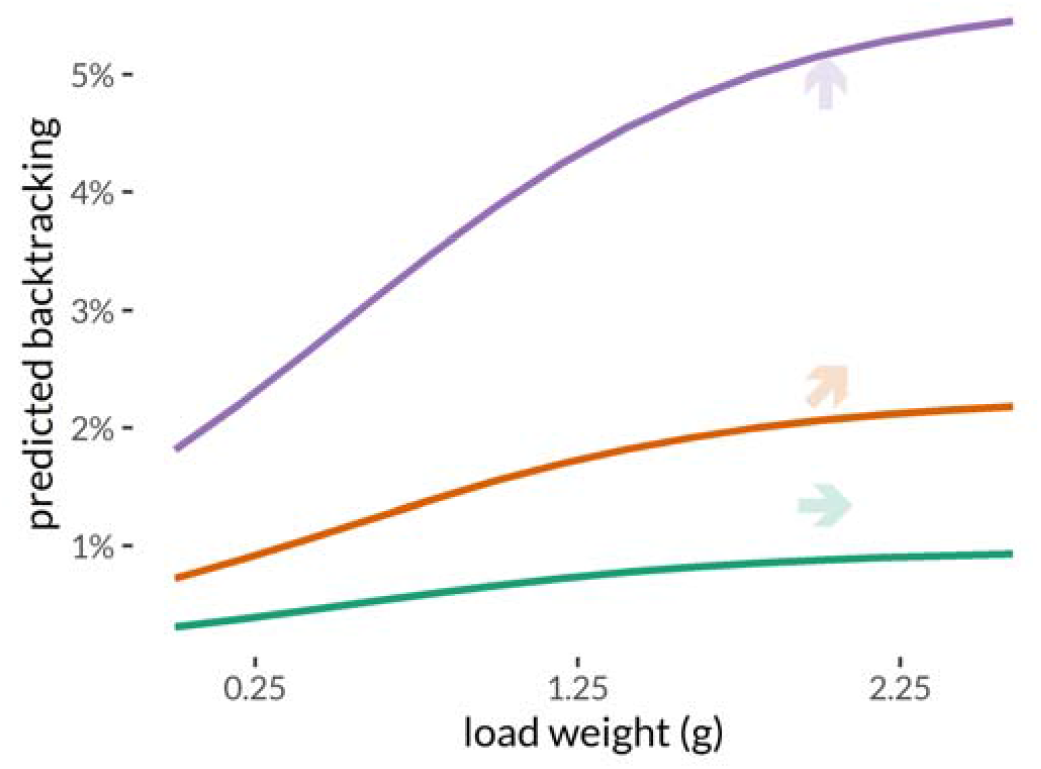
The percentage of backtracking predicted by the best fit mixed-effect hurdle model. Load weight predicts the probability that any backtracking occurs at all (lighter loads rarely slip), but once backtracking is happening, incline predicts how extreme it is (there is more severe backtracking on steeper slopes).

#### Path straightness

Teams took straighter paths when transporting lighter loads (* = -0.31, *p <* 0.001, *t*_*98*_ = -3.46) and when crossing less steep platforms (* = -0.30, *p <* 0.005, *t*_*98*_ = -3.37). However, we found no significant interaction between load weight and incline (Fig 6. *R*^*2*^ = 0.169). All things being equal, path straightness would be expected to decrease from ∼0.93 on a horizontal platform to ∼0.89 on a vertical surface and from ∼0.94 on a 0.25 gram load to ∼0.88 on a 2.25 gram load.

**Figure 6:**
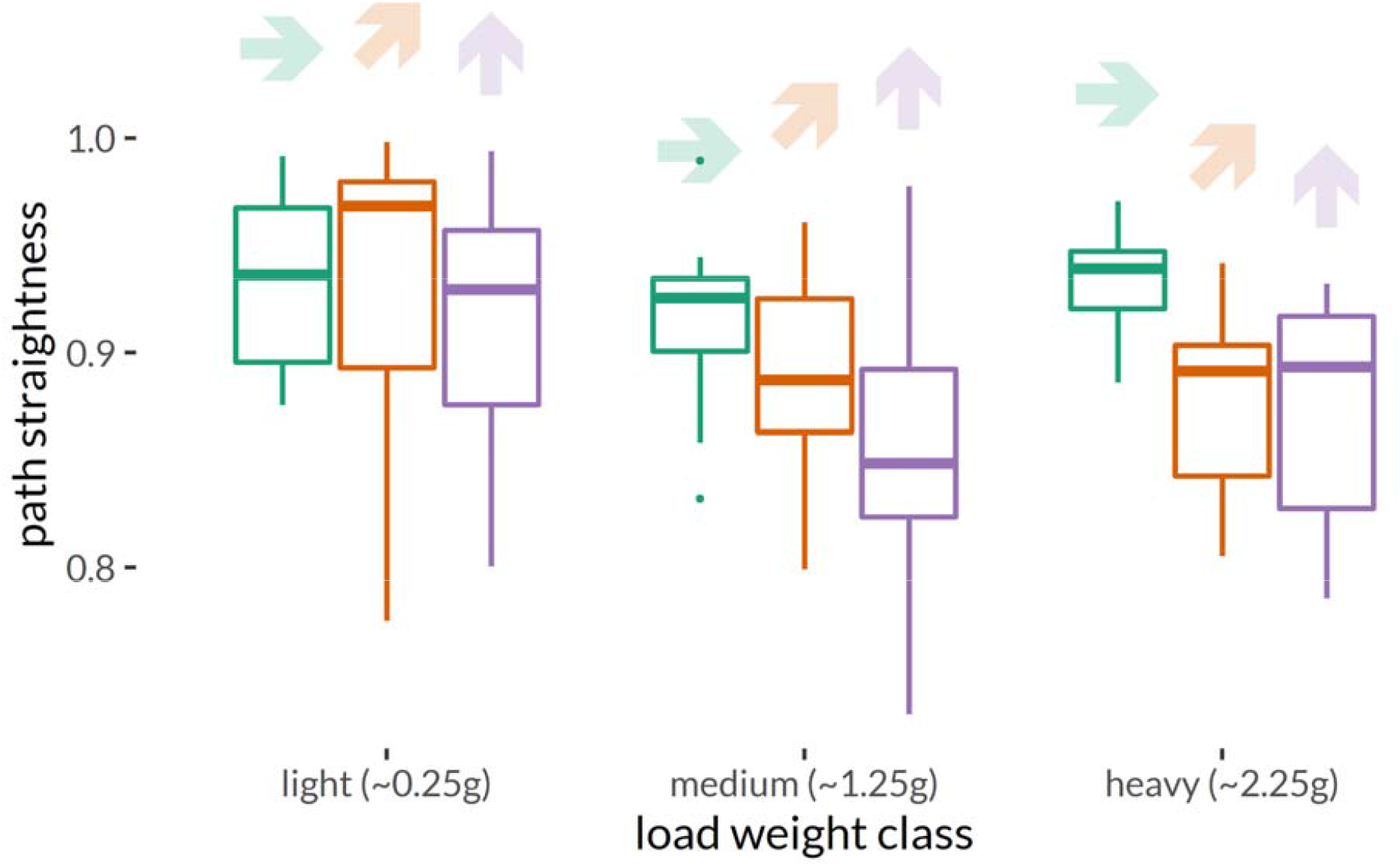
The path straightness of moving loads as a function of load weight and incline. Path straightness significantly decreases with heavier loads and with steeper inclines.

#### Instantaneous speed

To assess the speed teams can attain independent from the straightness of their path, we also calculated the average *instantaneous* speed—the speed of the load in any direction, moment-to-moment—excluding backtracking events. Ant teams moved *faster* on inclines (* = 0.06, *p* <0.005, *t*_*98*_ *=* 3.10) and slower while carrying heavier loads (* = -0.34, *p <* 0.001, *t*_*98*_ = -14.8), and we found no significant interaction between these factors (Fig 7. *R*^*2*^ = 0.661). However, because of the backtracking and longer paths taken on vertical surfaces, this increase in average instantaneous speed did not translate into a significantly faster average displacement speed from start to finish. All things being equal, average instantaneous speed would be expected to increase from ∼0.16 cm/s on a horizontal platform to ∼0.22 cm/s on a vertical surface and to decrease from ∼0.4 cm/s on a 0.25 gram load to ∼0.1 cm/s on a 2.25 gram load.

**Figure 7:**
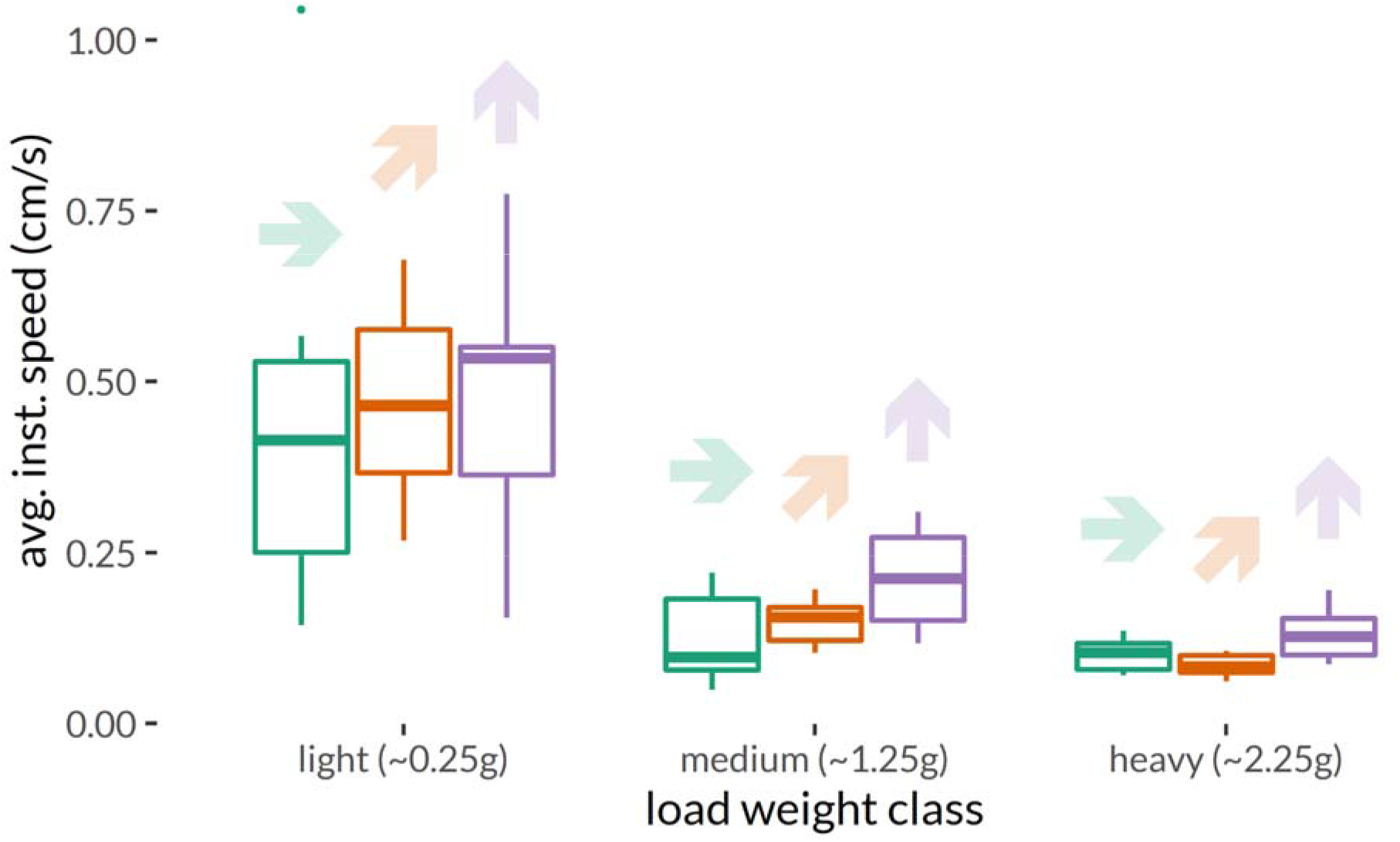
The average instantaneous load speed (excluding backtracking events) with respect to load weight and incline. The average instantaneous significantly decreases with heavier loads and significantly increases over steeper inclines.

### Quality versus quantity

The increase in instantaneous speed over steeper slopes could be explained by a change in the behavior of individual ants, a change in the group composition (e.g., a greater number of ants), or a combination of the two. Individual ants might remain engaged with the load longer on vertical surfaces. Theoretical and comparative work suggests that individual transporters’ persistence with loads can lead to greater success during collective transportation (McCreery et al. 2016, McCreery 2017), and the weaver ants might “hold on tighter” while on vertical surfaces, thus being better suited to maintain higher speeds. Although persistence is usually defined as a worker’s fidelity to a particular direction while pulling on an immobile load, in our case loads were being actively transported. Because the directional contribution of individual transporters could not be accurately estimated, we recorded how long they remained engaged with the load. Our measure of engagement is consequently analogous to a short term version of McCreery’s (2017) “total engagement effort” measure.

Alternatively (or in addition), changes in velocity may be due to changes in team size; the slight increase in group size on steeper slopes could explain the faster movement. Thus, we use measurements of ant persistence and group size to test both of these two hypotheses, which are not incompatible with each other.

#### Individual engagement with the load

We found that ants did not stay engaged with the load longer on steeper inclines (Fig 8A), suggesting that individual persistence is not one of the factors contributing to the observed increase in instantaneous speed on steeper slopes. However, the best model indicated that individual ants did remain engaged longer with lighter loads (* = 0.246, *p* <0.01, *z =* 2.8), which may suggest that among these ants, persistence is impeded by physical challenges as opposed to a response to them. We found a 35.4% increase in the expected probability of disengaging relative to a one gram increase in load weight (Fig 8B. *R*^*2*^ = 0.0399).

**Figure 8:**
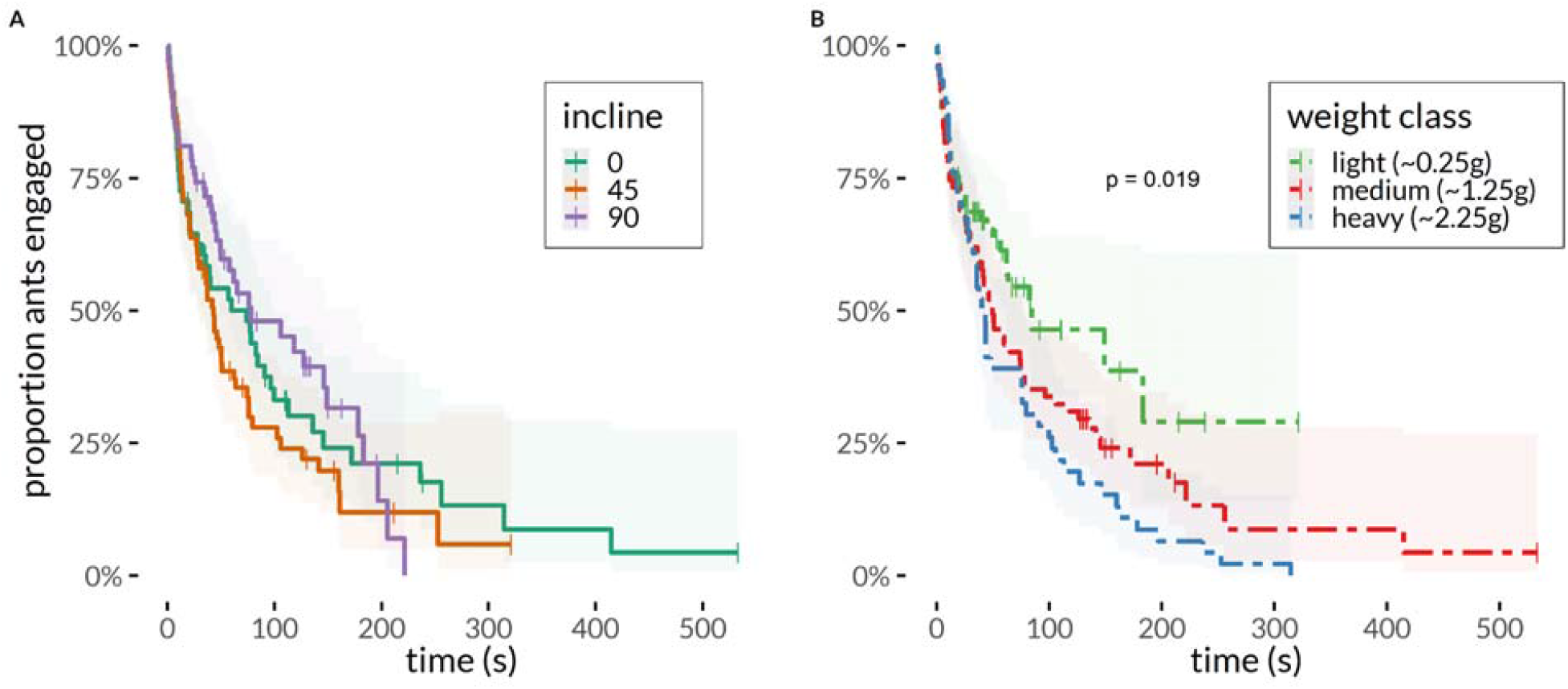
Duration that individual ants remain engaged with the load as a function of (A) incline and (B) load weight. (A) Note that incline does not affect load engagement, nor is it a significant predictor in any of the more complex models we examined. (B) Load weight did significantly predict how long ants remained engaged with loads; this panel represents the best fit model.

#### Team size and load speed

Our models indicated that the incline of the experimental platform had a significant statistical effect on the average instantaneous speed during transport. However, this may not represent an immediate, causal link between the two factors. Incline also affected the number of transporters engaged with the load. Thus, the number of transporters could potentially be acting as a mediator, and the incline would only indirectly affect the load’s speed via the number of ants involved.

As shown earlier, the regression of incline on the logged average instantaneous speed, ignoring the mediator (number of transporters), was significant, ( = 0.00330, *p <* 0.005, *t*_*98*_ *=* 3.10). Likewise, the regression of incline on the mediator variable (number of transporters), was also significant, ( = 0.0376, *p <* 0.001, *t*_*98*_ = 6.89). Additionally, the mediation process showed that the mediator variable had a significant effect on the logged average instantaneous speed, ( = 0.0446, *p <* 0.001, *t*_*97*_ = 2.40). Finally, the mediation analyses revealed that, after controlling for the mediator (number of transporters), incline was not a significant predictor of average instantaneous speed, ( = 0.00164, *p =* 0.199, *t*_*97*_ = 1.29).

We tested the significance of this indirect effect using quasi-Bayesian procedures. Unstandardized indirect effects were computed for each of 1,000 simulated samples, and the 95% confidence interval was computed by determining the indirect effects at the 2.5th and 97.5th percentiles. The bootstrapped unstandardized indirect effect was 0.00164, and the 95% confidence interval ranged from 0.000550 to 0.00348 (*p <* 0.001). Thus, we found that the number of transporters fully mediated the relationship between incline and average instantaneous speed (Fig 9).

**Figure 9:**
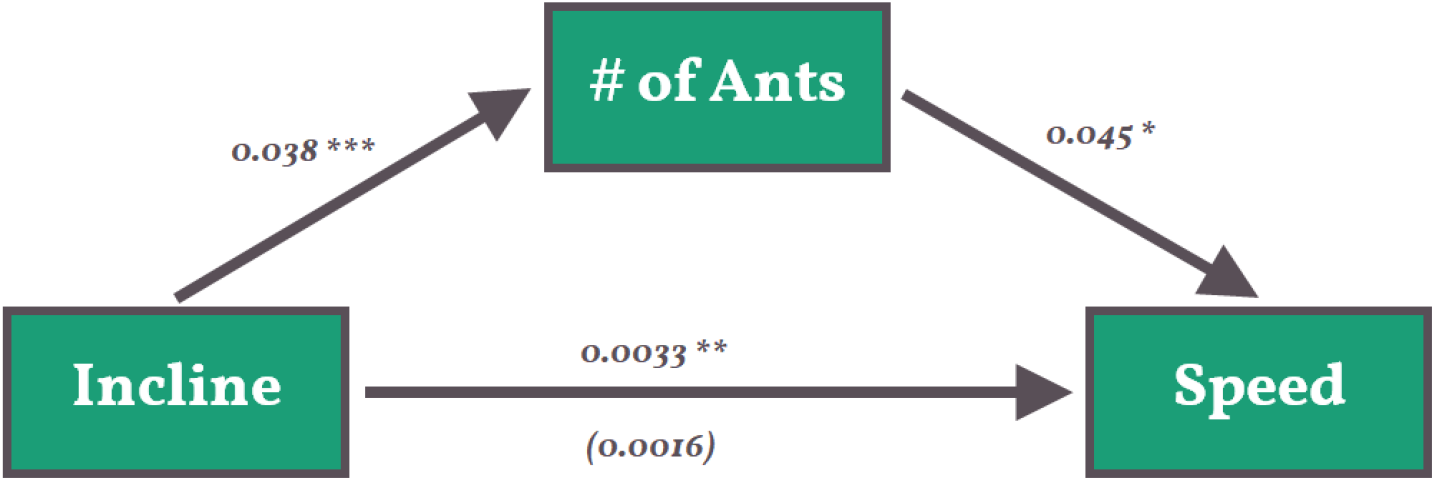
Mediation diagram depicting the unstandardized regression coefficients for relationships between platform inclination and the logged average instantaneous load speed, as mediated by the median number of transporters. Because incline’s effect on average instantaneous speed becomes insignificant after controlling for group size (the unstandardized regression coefficient is shown in parentheses), we can say that group size fully mediates the relationship.

## Discussion

We investigated how weaver ant transporter teams responded to loads of varying weights across inclines of varying angles. Our goal was to understand how transport efficiency (prey delivery rate or PDR) changed when teams were faced with these challenges. We also wanted to know how other more granular measures of group performance—like transport speed, path straightness, slipping and backtracking, and group size—interact to ultimately constitute the PDR.

### Adaptive allocation and prey delivery rate

Given previous studies on non-nomadic ant species, we expected that heavier loads (and likely steeper inclines) would decrease transport team efficiency. However, we found that weaver ant transport teams maintained a nearly constant per capita rate of return across our entire range of scenarios. From light loads carried over a flat plane to crickets weighing nine times as much and being pulled up a shear wall, the differences in PDR (per ant) were biologically negligible. This pattern of constant PDR can be visualized in the inversely proportional relationship between loads’ average displacement speed and their median weight per transporter (Fig 2B). Consequently, the higher the weight per ant (the “width” of the rectangle), the slower the load travels (the “height” of the rectangle). So a constant PDR indicates a possible fundamental trade-off between these two quantities and suggests a potential homeostatic regulatory mechanism for cooperative transport teams; ants are dynamically self-allocating to loads in a way that keeps the PDR constant. In other words, emergent regulation of constant PDR may be adaptive for colony foraging performance.

Constrained optimization provides one explanatory framework for the observed constant PDR. Assume that there are fitness benefits to a colony that can accumulate relatively large amounts of prey relatively quickly. This situation can be modeled with a fitness surrogate utility function:

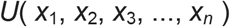

where there are *n* possible prey items to allocate ants to, *x*_*i*_ is the allocation of ants to prey item *i*, and *U* is the sum of all of their mass–delivery-speed products (where mass is a proxy for calories or a highly limited macronutrient, like protein). Fitness maximization would then be modeled as maximizing *U* over the different allocations of ants to prey items. However, because there are a finite number of ants, then all allocations must satisfy:

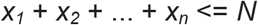

where *N* is the total number of foragers available to allocate. Alternatively, the problem could be posed as minimizing the number of ants *N* so long as the colony-level utility *U* meets a specific constraint. Constrained optimization problems like both of these can be solved with the method of Lagrange multipliers. Coincidentally, the solution to both formulations satisfies the condition that:

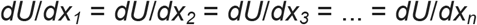

However, *dU/dx*_*i*_ is exactly the PDR associated with prey item *i*, and so the optimality condition is exactly a statement that the PDR should be equal across prey items at the optimal allocation. This result is also akin to the equilibrium solution of an ideal free distribution (IFD, (Fretwell and Lucas 1969) where the PDR plays the same role as patch suitability – ants allocate to different patches until there is no strong difference between suitability across those patches. However, in the IFD, the suitability functions represent proxies for individual rewards. Here, the individual-level suitability functions are the marginal returns of the colony-level utility, which is consistent with an inclusive-fitness perspective on IFD allocations. This result is also analogous to the Marginal Value Theorem (MVT) from Optimal Foraging Theory (OFT) (Charnov 1976, Stephens and Krebs 1987).

In other words, the PDR represents the marginal value of the next ant being added to a prey item, and the optimal allocation of a finite number of ants is the one in which the marginal values equalize across all items. In OFT, the focus is on how a single forager maximizes her rate of gain by allocating a minimal time budget across a portfolio of different patches. For the ants, a colony can allocate multiple individuals who focus on one patch (as opposed to one individual that moves among patches). However, the optimal result is the same – an individual should allocate itself to the patch with the highest marginal rate of return, and the allocation should stop when all individuals are allocated. In other words, this constant “rate of return” per ant is what one would expect if the colony were acting as an individual allocating its finite biomass (instead of its finite time) to different loads to optimize prey intake. Future research could test this hypothesis by comparing the PDR between very different colonies: although the optimality explanation is consistent with PDR regulation across prey items, it only predicts equalization within a colony, not between them. Colonies with different constraints (either in terms of nutrient constraints like prey availability or in terms of the number of available foragers) should differ in observed PDR.

### Constituent measures of group performance

We found that the challenge posed by steeper inclines qualitatively differed from that of increasing load weight, again contrary to our expectations. We also found no evidence of their interaction effects in our study. As with heavier loads, teams transporting over steeper slopes took more sinuous trajectories and had more ants engaged with the loads. But even though vertical surfaces were also associated with more backtracking (slipping), loads were transported up these slopes in the same amount of time. Interestingly, similar dynamics have been incidentally observed in the African species of weaver ant, *Oecophylla longinoda*. An unpublished study noted no difference between collective transport displacement speeds over horizontal and 35° inclines, though this result was tangential to the aim of their experiment (Lioni 2000).

### Faster average instantaneous speeds on vertical surfaces

Furthermore and counterintuitively, we found that loads were dragged significantly faster across vertical surfaces than over horizontal ones. We investigated two possible explanations by collecting further data from our recordings. Firstly, we hypothesized that the “effort” individual ants exert might be greater on vertical surfaces. McCreery (2017) suggested that longer engagement and persistence with loads correlates with more successful collective transport, thus individual weaver ants may stay more engaged in transport on steeper inclines. Our hypothesis predicts that ants will remain engaged with the load longer on vertical surfaces. Secondly, we hypothesized that the increased group sizes on vertical surfaces might lead to faster transport. Steeper slopes had larger team sizes, and larger team sizes might be the causal reason loads moved more quickly.

Our results suggest that the quantity of transporters (and not the persistence of individuals) is likely the direct reason for faster movement on vertical surfaces. A possible alternative explanation is that ant teams have less frictional forces to overcome on vertical surfaces; however the biomechanics of collective transport are more complicated than they might first appear. Ants climb by pulling *into* the walls (Endlein and Federle 2015), so any load being lifted by a tensile force will also experience a concomitant force pressing it into the wall. Regardless, it is not clear *how* this vertical condition leads to a larger number of ants in the transport team and *why* each team reaches the size that it does.

Our study raises many questions about the coordination of collective prey transport, both vertical and horizontal. Although the ultimate explanation for the weaver ants’ constant PDR might involve an optimized allocation of transport effort, the mechanisms behind this dynamic allocation are unclear. How are individual ants able to assess the marginal utility increase from engaging with a given load? We speculate that potential transporters may be able to measure the resistance and/or change in speed that results from an initial tug on a load. Previous studies have shown that some ant species will frequently over-recruit nestmates to pinned prey items, presumably because scouts are modulating recruitment based on the tractive resistance of the potential load (Hölldobler 1983, Detrain and Deneubourg 1997). We propose that a similar process may be used to assess loads during transport: if a moving load offers little resistance when pulled upon by a new ant, that potential transporter may be superfluous. Alternatively, the speed (or change in speed) of a load may also be important for this behavior. Swiftly moving, light loads may be less attractive to unengaged recruits while slower, heavier loads may be more attractive (even if workers disengage at higher rates from heavier loads).

### Future directions

This is the first study to investigate collective transport up vertical surfaces, and our findings suggest many potential avenues of further research, both for our system specifically and more generally across ant species. For example, the mere fact that transport efficiency did not decrease across any of our treatments suggests that the challenges given to these weaver ants were surmounted without detriment. It remains an open question: to what extent can PDR be regulated over a wider range of parameters? The “edge” of their ability to regulate is still unexplored.

Like previous studies of horizontal collective transport in weaver ants, we weighted our prey crickets with lead sinkers, placed the loads in high-traffic areas, and observed transport over relatively short distances (Lioni 2000). Our heaviest loads were much more dense than realistic prey items, and recent work has shown that load size and mass have different effects on transport in other ant species (McCreery et al. 2019). Our small but heavy crickets may have artificially limited the number of transporters that could easily engage with the load or, alternatively, decreased the usual friction for such a weight.

By placing the loads on pre-existing ant trails, we provided a steady stream of potential recruits that would not necessarily be present during normal prey retrieval. Although this choice decreased recruitment latency in our study, transport events further from the nest may exhibit different dynamics. Lastly, we only focused on the first ∼50 cm of collective transport. Since load speed over time is a nonlinear process (Lioni 2000) later portions of transport may unfold differently.

Lastly, there are other types of inclined surfaces that we did not test. Weaver ants will routinely transport prey along the *underside* of surfaces, and performance could change whether the teams are transporting loads up or down an incline. We also did not test how teams respond to transitions between horizontal and inclined surfaces: such transitions are common in nature and might pose challenges in and of themselves.

## Funding

This work was supported by the Australian Department of Education and Training’s Endeavour Research Fellowship and the National Science Foundation’s Graduate Research Opportunities Worldwide.

## Acknowledgements

We would like to thank Kelly S. O’Meara for her help in recording the transport events and the support she offered in the field.

## Notes

### Competing Interest Statement

The authors have declared no competing interest.

